# Scaling of Noise Under Resource Constraints in Gene Regulatory Motifs

**DOI:** 10.64898/2026.08.02.742368

**Authors:** Utkarsh Singh Solanki, Abhilash Patel, Abhyudai Singh

## Abstract

Understanding noise propagation in gene regulatory circuits requires accounting for both model and resource constraints. In this work, we investigated the role of model order in influencing stochastic behaviour by deriving and analytically comparing reduced protein-only models with higher-order models that include mRNA and molecular complexes, and found that protein-based models can exhibit higher noise levels in the gene expression. Through frequency-response analysis, we explained that the higher-order models provide additional noise-filtering effects. We also analyzed a one-dimensional constrained model and showed that the Fano factor decreases as the strength of resource constraint increases. Finally, we considered larger circuit motifs, such as toggle switches and incoherent feed-forward loops, and found that resource limitations can minimise stochastic switching in a bistable circuit, whereas in an incoherent feed-forward loop, resource constraints can make the adaptation faster. Our results highlight that both mechanistic detail and shared resource constraints play a central role in determining fluctuation levels in biomolecular circuits.

## I. Introduction

In synthetic biology, elementary biochemical reactions are configured to achieve a desired phenotype in the cell by altering gene expression, a stochastic process in which genes are transcribed and translated into mRNAs and proteins [1]–[3]. Due to the inherent stochastic fluctuations in gene expression, designing predictable circuit behaviour is a fundamental challenge [4], [5]. There can be various noise sources contributing to the variability of the circuit, including mRNA birth-death fluctuations, transcriptional and translational bursting [6], and limited abundance of resources such as ribosomes and RNA polymerases [7]–[10]. Despite identical inputs and environmental conditions, these sources of stochasticity can lead to significant variability in the desired output of a biomolecular system [11]–[22].

Stochastic analysis of gene expression has been considerably studied in the literature. Prior work on resource-aware modeling [23], [24], the effects of ribosomal constraints on stochastic gene expression [25], and the impact of resource scarcity on toggle switches [26] provides a foundation for analyzing gene expression under shared cellular resources. These contributions offer deep insight into deterministic and stochastic resource-aware modeling. However, a systematic examination of how noise propagates across models of different complexity, combined with an extended study of resource effects on gene regulatory motifs, has not been performed.

The limited availability of cellular resources, such as RNA polymerases and ribosomes, which are essential for gene expression and protein synthesis, creates a competitive environment within the cell [27]. Among many cellular resources, ribosomes play a pivotal role by binding to mRNA molecules to form translation complexes. When multiple genes are expressed simultaneously, or when multiple mRNAs from the same gene compete for a shared finite pool of ribosomes, unintended coupling between genetic components emerges, leading to deviations from expected circuit behavior [28], [29]. These limitations influence the stochastic nature of gene expression and affect the variability, robustness, and overall reliability of circuit performance. The critical impact of finite cellular resources on the stochastic dynamics of gene regulatory networks therefore makes it essential to integrate resource-aware considerations into the design and modeling of biomolecular systems.

In this work, we investigate the effects of resource constraints on noise propagation in biomolecular systems. We first obtain reduced-order models that incorporate the availability of free ribosomes as a constraint. Using stochastic simulations, we study the effect of constraints on gene expression dynamics by computing the Fano factor (variance-to-mean ratio). We then evaluate how well the reduced-order models capture resource constraint effects by comparing them against the full-order model. Through frequency-response analysis, we find that protein-only reduced models exhibit higher noise levels, whereas full-order models with intermediate species provide additional noise filtering. Finally, we extend our analysis to complex gene regulatory motifs, including the bistable toggle switch and the incoherent feedforward loop. Our results reveal that resource constraints reduce stochastic switching frequency in bistable toggle switches and lower the Fano factor in incoherent feedforward loops while preserving adaptation capability.

## II. Model Formulation

We consider a model of constitutive gene expression in which multiple mRNAs from a single gene compete for a shared, finite pool of ribosomes. Starting from a three-dimensional (3D) stochastic model that explicitly captures mRNA, ribosome–mRNA complex, and protein copy numbers, which is reduced to a 2D model capturing mRNA and protein copy numbers, which is further reduced to a 1D Model capturing only protein copy numbers.

### A. 3D Model

The biochemical reactions governing gene expression with explicit ribosome binding are

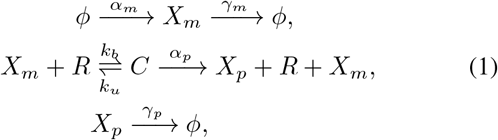

where *X*_*m*_ denotes mRNA, *C* the ribosome–mRNA translation complex, *X*_*p*_ the protein, and *R* the free ribosome. We assume that ribosome binding protects mRNA from degradation and that the total ribosome count *r*_*T*_ is conserved, so the free ribosome level satisfies *r* = *r*_*T*_ − ***c***.

The stochastic model is defined by the propensity functions, that determines the probability of occurrence of a reaction per unit time in Table II, which govern the time evolution of the integer-valued copy numbers ***x***_***m***_, ***c, x***_***p***_ ∈ {0, 1, … }, where bold lower case notation is used to denote species counts, indicating random processes. The mean values at steady state [25], [30] can be given as:

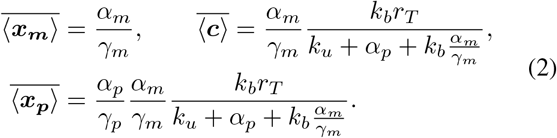

where, 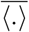 represents the mean value at steady state and with the limiting case of fast translation, the mean protein count at steady state simplifies to

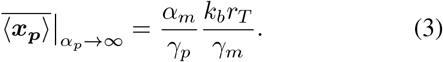

**TABLE I.**
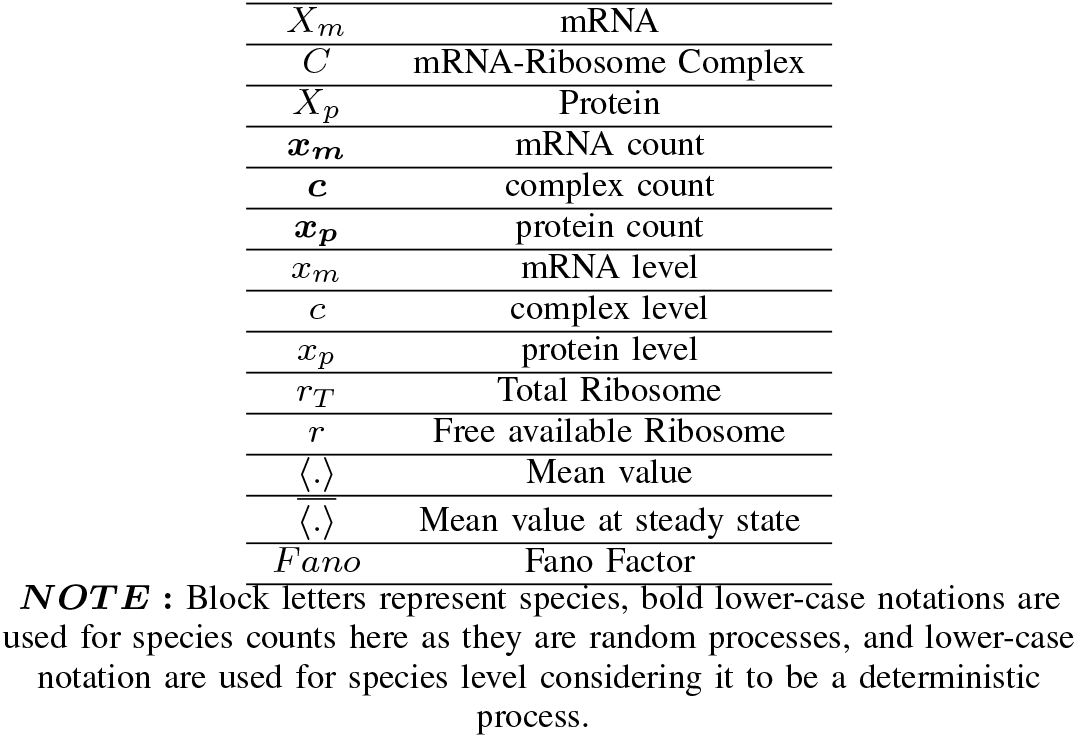
A SUMMARY OF THE NOTATION USED.

**TABLE II.** Propensities for population reset for corresponding reactions for single gene expression 3D MODEL.

| Reaction | Population Reset | Propensity |
| --- | --- | --- |
| $\phi \rightarrow X_m$ | $\mathbf{x}_m \rightarrow \mathbf{x}_m + 1$ | $\alpha_m$ |
| $X_m \rightarrow \phi$ | $\mathbf{x}_m \rightarrow \mathbf{x}_m - 1$ | $\gamma_m \mathbf{x}_m$ |
| $X_m + R \rightarrow C$ | $\mathbf{c} \rightarrow \mathbf{c} + 1,$<br>$\mathbf{x}_m \rightarrow \mathbf{x}_m - 1$ | $k_b (r_T - \mathbf{c}) \mathbf{x}_m$ |
| $C \rightarrow X_m + R$ | $\mathbf{c} \rightarrow \mathbf{c} - 1,$<br>$\mathbf{x}_m \rightarrow \mathbf{x}_m + 1$ | $k_u \mathbf{c}$ |
| $C \rightarrow X_p + X_m + R$ | $\mathbf{x}_p \rightarrow \mathbf{x}_p + 1,$<br>$\mathbf{c} \rightarrow \mathbf{c} - 1,$<br>$\mathbf{x}_m \rightarrow \mathbf{x}_m + 1$ | $\alpha_p \mathbf{c}$ |
| $X_p \rightarrow \phi$ | $\mathbf{x}_p \rightarrow \mathbf{x}_p - 1$ | $\gamma_p \mathbf{x}_p$ |

The stochasticity in protein expression can be quantified using the Fano factor, widely used to quantify noise in gene expression, defined as the variance-to-mean ratio of protein copy numbers [31], [32]. The steady-state Fano factor of the protein copy number obtained from Linear Noise Approximation (LNA) [33]–[36] is Eq. (4) (see the top of the following page), and in the limiting cases, the Fano factor simplifies to,

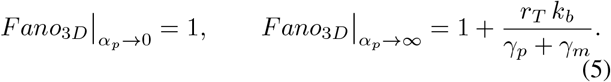

The lower limit (*α*_*p*_ → 0), recovers Poissonian statistics (no translational bursting), while the fast translation (*α*_*p*_ → ∞), shows that noise grows with the total ribosome count *r*_*T*_ and recovers the Fano factor as obtained in the classical model of stochastic gene expression with no constraints on ribosome availability [14], [37].

For the fixed mean protein count, from Eq. (2), we have,

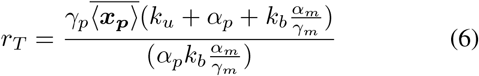

and with the limiting cases of translation rate, the Fano factor simplifies to,

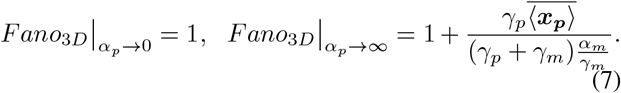

### B. Reduced 2D Model

When ribosome binding and unbinding are fast relative to mRNA and protein turnover (*k*_*b*_, *k*_*u*_ ≫ *γ*_*m*_, *γ*_*p*_), the complex ***c*** equilibrates rapidly given the current mRNA level. Eliminating the complex via a quasi-steady-state approximation yields a reduced stochastic process on the state (***x***_***m***_, ***x***_***p***_) ∈ {0, 1, …} with the propensities in Table III.

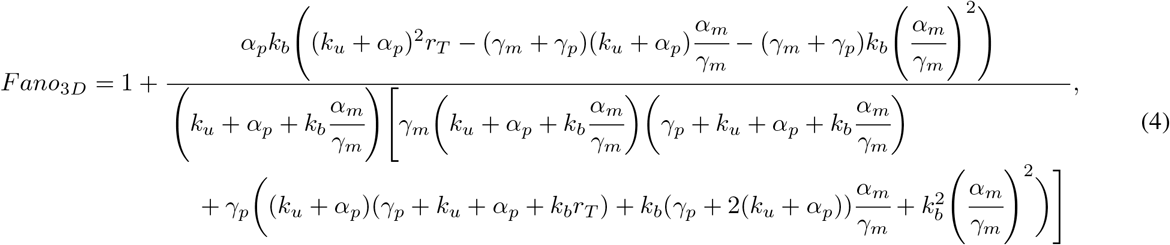

**TABLE III.**
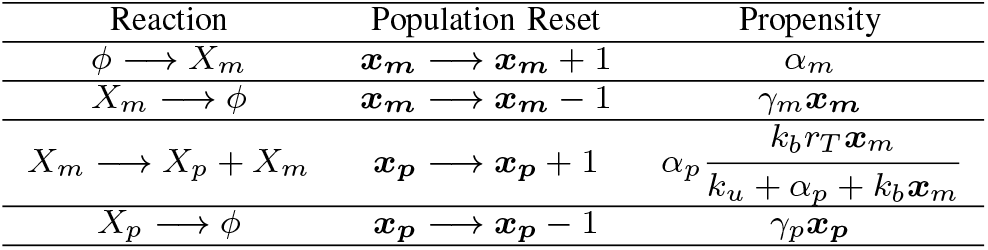
Propensities for population reset for corresponding reactions for single gene expression 2D MODEL.

The key consequence is that the protein production propensity becomes a nonlinear, saturating function of the mRNA count:

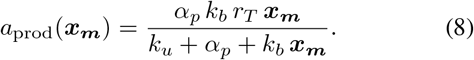

This is a Michaelis–Menten-like rate in the mRNA copy number, with the effective dissociation constant being (*k*_*u*_ + *α*_*p*_)*/k*_*b*_. At low mRNA counts (*k*_*b*_ ***x***_***m***_ ≪ *k*_*u*_ + *α*_*p*_), the rate is approximately linear in ***x***_***m***_ and the system behaves as if resources are abundant. At high mRNA counts, the rate saturates at *α*_*p*_*r*_*T*_, reflecting complete ribosome utilization.

We have used these two models explicitly in our previous work [25] to investigate stochastic gene expression.

## III. 1D Model

In order to reduce the order of a simple gene expression model to obtain a protein-only one-dimensional (1D model), we undergo an approximation that the mRNA half-life is significantly less than the protein half-life, *γ*_*m*_ >> *γ*_*p*_, such that proteins are produced in bursts. Using this approximation, we can further ignore the mRNA dynamics and obtain a model where protein synthesis occurs as a bursty birth-death process, where proteins are produced in instantaneous bursts at a rate *α*_*m*_, with each burst generating *B* molecules, where *B* is an independent and identically distributed random variable.

For this bursty birth-death process, the steady-state statistics of the mean protein level and its Fano factor are given by

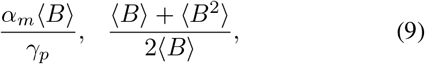

respectively [32], [38], [39], where ⟨.⟩ represents the expected value. Note that *B* = 1 with probability one (non-bursty synthesis) reduces the Fano factor to one in Eq. (9) corresponding to Poissonian copy-number fluctuations.

Comparing the steady-state mean protein level reported in Eq. (2) with the mean in Eq. (9) yields the mean burst size as

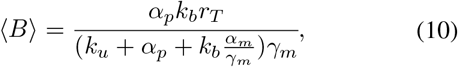

Taking the limit *γ*_*p*_ → 0 (i.e., protein dynamics is much slower compared to its mRNA), the Fano factor in Eq. (4) reported for the full 3D model reduces to

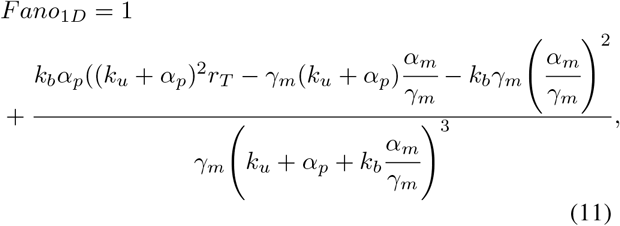

Comparing *Fano*_1*D*_ to the Fano factor for a bursty birth-death process (second term in Eq. (9)) results in the second-order statistical moment of *B* given by:

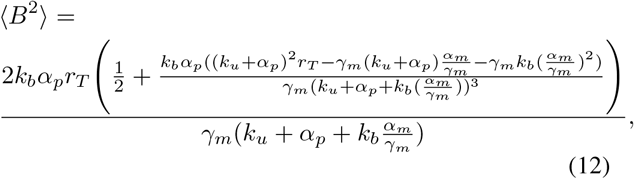

Thus, our one-dimensional model of gene expression with resource competition reduces to a bursty birth-death process (Table IV), with burst statistics given by Eqs. (10) and (12). For the purpose of simulations, we use negative binomially distributed bursts with these statistics. In the limit of fast translation *α*_*p*_ → ∞ (no resource constraints), this model reduces to the classical one-dimensional model of gene expression with geometrically distributed bursts with mean burst size [40]

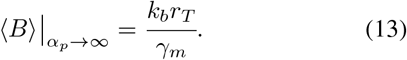

**TABLE IV.** Propensities for population reset for corresponding reactions for single gene expression 1D MODEL.

| Reaction | Population Reset | Propensity |
| --- | --- | --- |
| $\phi \rightarrow X_p$ | $\mathbf{x}_p \rightarrow \mathbf{x}_p + B$ | $\alpha_m$ |
| $X_p \rightarrow \phi$ | $\mathbf{x}_p \rightarrow \mathbf{x}_p - 1$ | $\gamma_p \mathbf{x}_p$ |

## IV. Simulations comparison

We compared the protein levels and the corresponding Fano factor for full order (3D model, TABLE II) and reduced order models, comprising of mRNA and protein dynamics (2D model, TABLE III) and protein only dynamics (1D model, TABLE IV). We performed stochastic simulations using the Gillespie algorithm [41], [42] for all the models, establishing the common observational fact that resource constraint leads to decrease in mean protein levels as well as decrease in the variability in the mean protein levels due to increased ribosome interaction and its finite availability. In the scenario, with abundance of available resources (i.e. higher translation rate), the mean protein levels for all the models overlapped each other (Fig. 3(a)) with Fano Factor also overlapping but slightly comparable in the order, 1D > 2D > 3D (Fig. 3(b)).

**Fig. 1.**
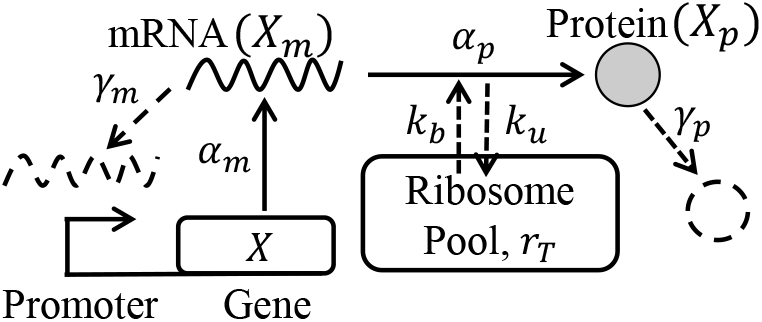
Schematic for simple gene expression with a finite ribosome pool. Gene *X* transcribes mRNA *X*_*m*_ at *α*_*m*_, while the mRNA is degraded and diluted effectively at a rate *γ*_*m*_. The mRNA *X*_*m*_ binds to ribosomes from a finite pool *r*_*T*_ at a rate *k*_*b*_ and dissociates at a rate *k*_*u*_. The ribosome-mRNA complex translates protein *X*_*p*_ at a rate *α*_*p*_, which subsequently undergoes dilution and degradation at an effective rate *γ*_*p*_.

**Fig. 2.**
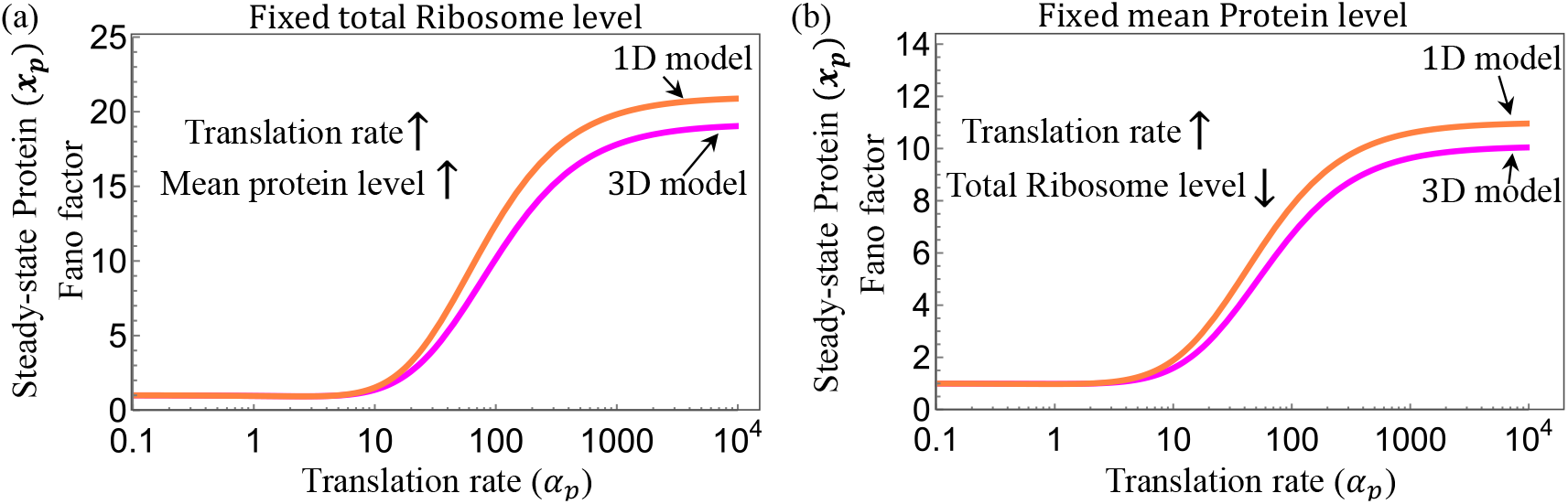
The steady-state Fano factor for quantifying fluctuations in the protein counts in the full model (Eq. 4) vs the reduced model (Eq. 11). The key parameters used are: *α*_*m*_ = 10, *γ*_*m*_ = 10, *γ*_*p*_ = 1, *k*_*u*_ = 0, *k*_*b*_ = 20. (a) for fixed total ribosome level (*r*_*T*_ = 10), (b) for fixed mean protein level 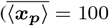 in Eq. (6)).

**Fig. 3.**
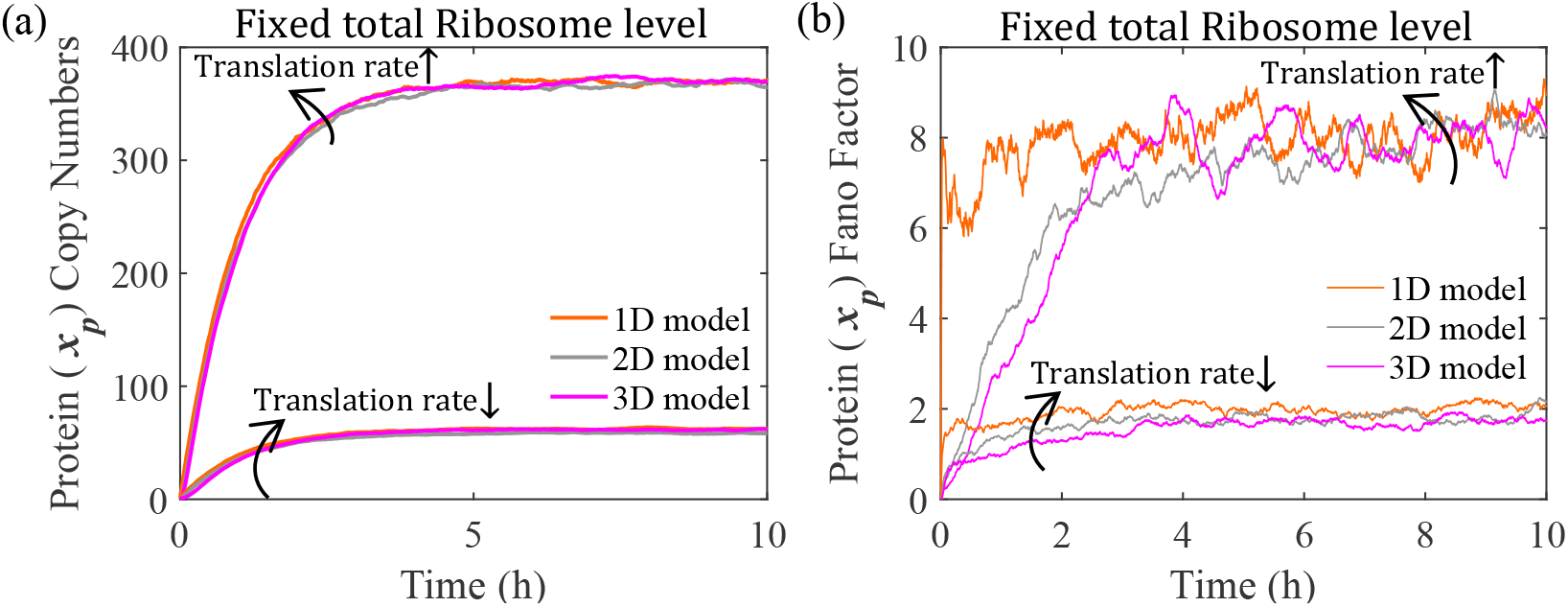
Simulations for simple gene expression, comparing all models in Table II, III, IV. The key parameters for the model used were considered as follows: *r*_*T*_ = 50 molecules per cell, *k*_*b*_ = 5 *h*^−1^, *k*_*u*_ = 50 *h*^−1^, *γ*_*m*_ = 30 *h*^−1^, *α*_*m*_ = 50 *h*^−1^, and *γ*_*p*_ = 1 *h*^−1^. All these parameters are scaled with respect to the protein decay rate. The simulation was performed for 500 trajectories. (a) An increase in the translation rate increases the mean protein copy numbers in a scenario without resource limitation (*α*_*p*_ = 400) and under resource limitation (*α*_*p*_ = 10) the mean protein copy number of all three models decreases with similar mean levels of protein copy numbers. (b) Higher Fano factor, variance over mean for 500 trajectories, of ***x***_*p*_ without resource limitation (*α*_*p*_ = 400) and under resource limitation (*α*_*p*_ = 10) Fano factor decreases for all the models.

To further examine the effect of reducing the model order, we observe the mean protein levels and their corresponding Fano Factor under limited resources (i.e. lower translation rate), where, similar to the resource abundance scenario, the mean protein levels overlapped each other for all the models (Fig. 3(a)) with Fano Factor also overlapping but slightly comparable in the order, 1D > 2D > 3D (Fig. 3(b)). Although the Fano factor tends to overlap for corresponding parameters in Fig. 3, but do not exactly replicate each other. This leads to the observation that, in general, protein-only models can exhibit higher noise levels, while intermediate species in detailed models provide additional noise-filtering effects.

### A. Frequency-response analysis

The observation that protein-based models can exhibit higher noise levels, while intermediate species in detailed models provide additional noise-filtering effects, can be validated using the frequency response analysis.

In a deterministic framework, the set of chemical reactions in Eq. (1) can be modeled, using the mass action law, as

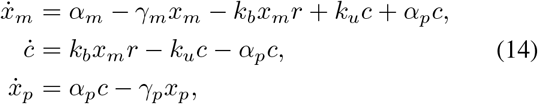

where, *x*_*m*_, *c, x*_*p*_, *r* represent concentrations of *X*_*m*_, *C, X*_*p*_ and resources *R* respectively. To capture the effects of resource constraint, we define the total finite cellular resources *r*_*T*_ as the sum of freely available resources *r* and ribosomes bound to mRNA molecules in the form of translation complex *c*, as derived in the previous work [25], Under the assumption, that there finite resources, *r* = *r*_*T*_ − *c*, the Eq. (14) can be written as,

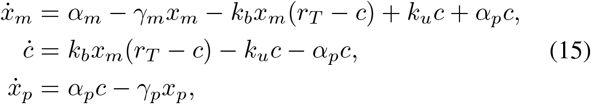

We get the non-linear model as:

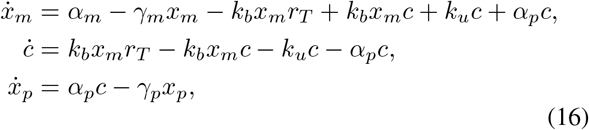

Taking, input, *u* = *α*_*m*_, and output, *y* = *x*_*p*_, Computing transfer function, upon linearization around equilibrium points,

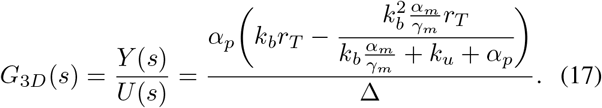

where the Δ is

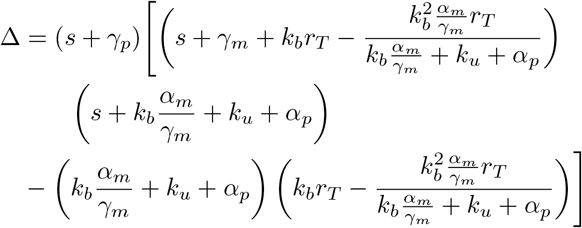

Under the assumption of quasi-steady state approximation for the dynamics of the complex *c* (*ċ* = 0), the model reduces to two-dimensional as:

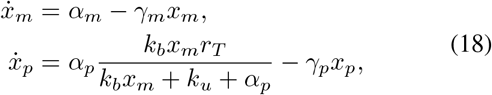

Therefore, the transfer function becomes

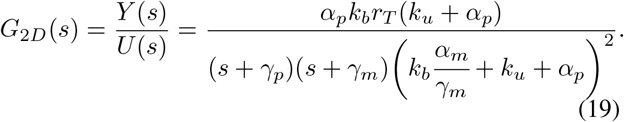

Under the assumption of quasi-steady state approximation for the dynamics of the mRNA, 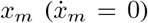, the model reduces to one-dimensional:

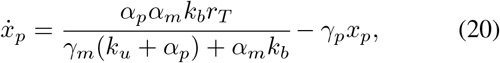

Therefore, the transfer function becomes,

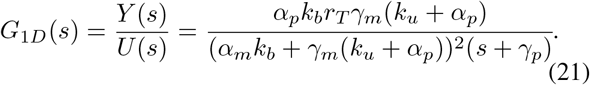

Using, transfer functions for all the models, in Eqs. (17), (19) and (21), we plot the Bode magnitude plot (Fig. 4), to analyze the system behavior. The gain plot indicates the same low-frequency gain for all the models, and for high-frequency regions, we use the notion of filtering, which indicates how well the system attenuates high-frequency noise.

**Fig. 4.**
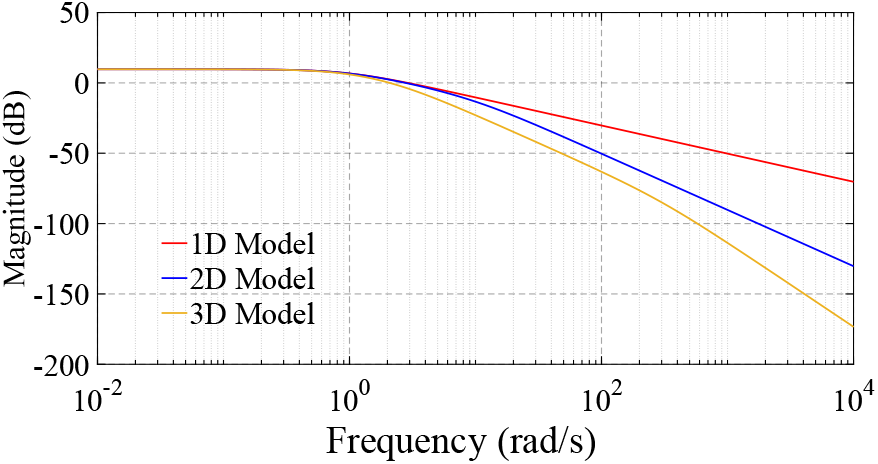
Bode magnitude plot for 1D, 2D and 3D models of simple gene expression with transfer functions in Eqs. 17, 19, 21 with resource limitation (*α*_*p*_ = 10) for input, *u* = *α*_*m*_, and output, *y* = *x*_*p*_ . The response shows a higher roll-off rate for the 3D model compared to the 1D model. The key parameters for the model used were considered as follows: total ribosome numbers assumed to be *r*_*T*_ = 50 molecules per cell and rates *k*_*b*_ = 5 *h*^−1^, *k*_*u*_ = 50 *h*^−1^, *γ*_*m*_ = 10 *h*^−1^, *α*_*m*_ = 20 *h*^−1^, *α*_*p*_ = 10 *h*^−1^ and *γ*_*p*_ = 1 *h*^−1^.

Firstly, the Bode magnitude plot for the 1D model shows a slope of −20 dB/decade, representing a single pole, representing decrease in gain with increase in frequency, leading to the system attenuating high-frequency noise and allowing low-frequency signals to pass. In the 2D model, a slope of −40 dB/decade indicates the presence of two poles, making the gain drop further and leading to stronger suppression of high-frequency noise. Similarly, in the 3D model, we observe a slope of −60 dB/decade corresponding to the three poles, where the gain further drops and the system strongly filters out high-frequency noise.

## V. Effect of Resource on Noise propagation in Complex Biomolecular Circuits

To investigate the impact of resource constraints on larger gene regulatory networks, we consider two complex circuits, the bistable toggle switch [43] and incoherent feedforward loop [44].

### A. Bistable toggle switch

In a toggle switch, gene *X* and gene *Y* encoding repressor proteins *X*_*p*_ and *Y*_*p*_, repress each other (Fig. 5(a), inset). A toggle switch is a bistable system, having two stable steady states, and noise can switch the states in between these, irrespective of the initial condition. The propensity of underlying reactions in the toggle switch is presented in Table V. The mRNAs *X*_*m*_ and *Y*_*m*_ are transcribed with the propensities *α*_*mx*_*f* (***y***_***p***_) and *α*_*my*_*g*(***x***_***p***_), where, 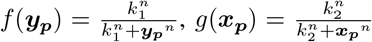, with *k*_1_ and *k*_2_ as the dissociation constant of *Y*_*p*_ and *X*_*p*_, while mRNA *X*_*m*_ and *Y*_*m*_ are degraded and diluted effectively with the propensity *γ*_*mx*_***x***_***m***_ and *γ*_*my*_***y***_***m***_. The ribosome binds to mRNA *X*_*m*_ with propensity *k*_*bx*_***x***_***m***_(*r*_*T*_ ***c***_***x***_ ***c***_***y***_) and unbinds with propensity *k*_*ux*_***c***_***x***_. Similarly, ribosome binds and unbinds to mRNA *Y*_*m*_ with propensity *k*_*by*_***y***_***m***_(*r*_*T*_ − ***c***_***x***_ − ***c***_***y***_) and unbinds with the propensity *k*_*uy*_***c***_***y***_, where *r*_*T*_ represents the pool of ribosomes. The proteins *X*_*p*_ and *Y*_*p*_ are synthesized with the propensities *α*_*px*_***c***_***x***_ and *α*_*py*_***c***_***y***_ respectively. Then, the proteins degrade with the propensities *γ*_*px*_***x***_***p***_ and *γ*_*py*_***y***_***p***_ respectively.

**TABLE V.** Propensities for population reset for corresponding reactions for Toggle switch.

| Reaction | Population Reset | Propensity |
| --- | --- | --- |
| $\phi \xrightarrow{f(y_p)} X_m$ | $x_m \rightarrow x_m + 1$ | $\alpha_{mx} f(y_p)$ |
| $X_m \rightarrow \phi$ | $x_m \rightarrow x_m - 1$ | $\gamma_{mx} x_m$ |
| $X_m + R \rightarrow C_X$ | $c_x \rightarrow c_x + 1,$<br>$x_m \rightarrow x_m - 1$ | $k_{bx} x_m (r_T - c_x - c_y)$ |
| $C_X \rightarrow X_m + R$ | $c_x \rightarrow c_x - 1,$<br>$x_m \rightarrow x_m + 1$ | $k_{ux} c_x$ |
| $C_X \rightarrow X_p + X_m + R$ | $x_p \rightarrow x_p + 1,$<br>$c_x \rightarrow c_x - 1,$<br>$x_m \rightarrow x_m + 1$ | $\alpha_{px} c_x$ |
| $X_p \rightarrow \phi$ | $x_p \rightarrow x_p - 1$ | $\gamma_{px} x_p$ |
| $\phi \xrightarrow{g(x_p)} Y_m$ | $y_m \rightarrow y_m + 1$ | $\alpha_{my} g(x_p)$ |
| $Y_m \rightarrow \phi$ | $y_m \rightarrow y_m - 1$ | $\gamma_{my} y_m$ |
| $Y_m + R \rightarrow C_Y$ | $c_y \rightarrow c_y + 1,$<br>$y_m \rightarrow y_m - 1$ | $k_{by} y_m (r_T - c_x - c_y)$ |
| $C_Y \rightarrow Y_m + R$ | $c_y \rightarrow c_y - 1,$<br>$y_m \rightarrow y_m + 1$ | $k_{uy} c_y$ |
| $C_Y \rightarrow Y_p + Y_m + R$ | $y_p \rightarrow y_p + 1,$<br>$c_y \rightarrow c_y - 1,$<br>$y_m \rightarrow y_m + 1$ | $\alpha_{py} c_y$ |
| $Y_p \rightarrow \phi$ | $y_p \rightarrow y_p - 1$ | $\gamma_{py} y_p$ |

$f(y_p) = \frac{k_1^n}{k_1^n + y_p^n}$ , $g(x_p) = \frac{k_2^n}{k_2^n + x_p^n}$ are repressing functions with $n = 2$ . They scale the transcription rate.

**Fig. 5.**
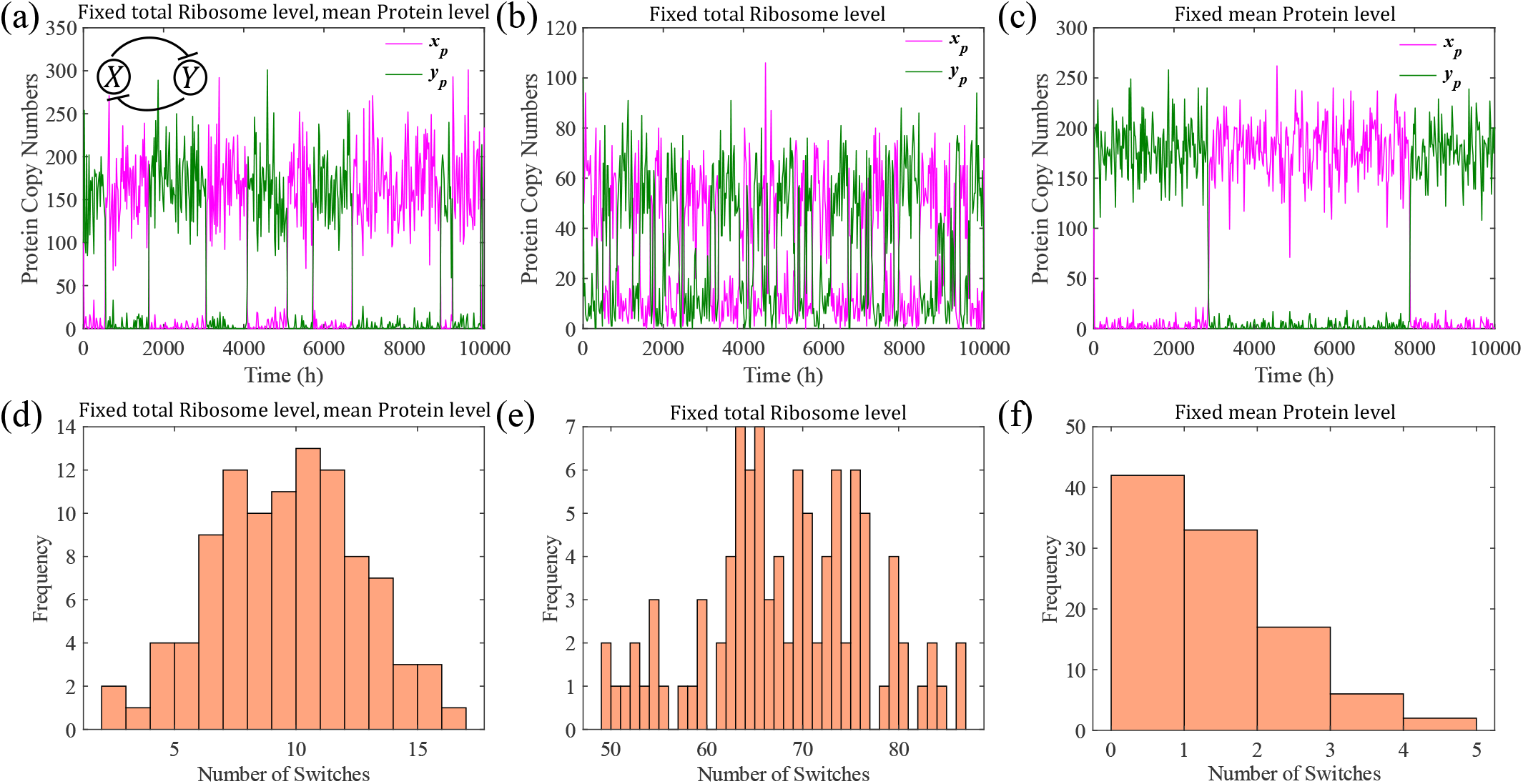
Simulations for Toggle switch (inset, a) (Table V). The key parameters for the model used were considered as follows: the rates are *α*_*mx*_ = *α*_*my*_ = 10 *h*^−1^, *γ*_*mx*_ = *γ*_*my*_ = 1 *h*^−1^, *k*_1_ = *k*_2_ = 20, *k*_*bx*_ = *k*_*by*_ = 1 *h*^−1^, *k*_*ux*_ = *k*_*uy*_ = 10 *h*^−1^, *γ*_*px*_ = *γ*_*py*_= 1 *h*^−1^. All these parameters are scaled with respect to the protein decay rate. The simulation was performed for 100 trajectories with cooperativity as *n* = 2. (a) Sample trajectory of ***x***_*p*_ and ***y***_*p*_ counts in a scenario without resource limitation due to *α*_*p*_ = 80 for 100 trajectories, and the total ribosome per cell *r*_*T*_ is given as 20 molecules. (b) Sample trajectory of ***x***_*p*_ and ***y***_*p*_ counts in a scenario with resource constraint due to *α*_*p*_ = 10 for 100 trajectories, and the total ribosome per cell *r*_*T*_ is given as 20 molecules. (c) Sample trajectory of ***x***_*p*_ and ***y***_*p*_ counts in a scenario with constraint due to *α*_*p*_ = 10 for 100 trajectories, and the increased total ribosome per cell *r*_*T*_ given as 55 molecules, to maintain the protein copy numbers the same as without resource constraint. The Histogram represents the distribution of the number of switches between ***x***_*p*_ and ***y***_*p*_ per trajectory for 100 trajectories in a scenario (d) without resource limitation, (e) with resource limitation, (f) with the mean protein copy numbers fixed and limited resources.

This toggle switch is simulated using the Gillespie algorithm, as done in previous sections, to capture the noise propagation. The simulation is performed for 100 trajectories, and the noise-induced switching frequency is obtained. We considered two scenarios. The first is of the total ribosomes fixed, and the translation rate *α*_*px*_ = *α*_*py*_ = *α*_*p*_ is varied to capture resource constraints. This reduces the mean protein count of the ‘higher’ stable steady-state. The second scenario ensures that the mean protein copy number is fixed; hence, we vary the total ribosomes *r*_*T*_ in proportion to the change in the translation rate *α*_*p*_. A sample trajectory for protein copy number for the toggle switch is shown in (Fig. 5(a)) for an unconstrained scenario, where *α*_*p*_ = 80. To induce constraint, we changed *α*_*p*_ to 10. In Fig. 5(b), a sample trajectory for a scenario of fixed total ribosome *r*_*T*_ is presented. We can note that the stable steady level decreased, with significantly large switching. In contrast to this, in the scenario where total ribosome *r*_*T*_ is varied alongside the translation rate *α*_*p*_ = 10, it showed less switching in addition to maintaining the unconstrained-level steady state (Fig. 5(c)).

We quantified the number of switches for each trajectory, and the histogram plots showed how frequently two system states, protein *X*_*p*_ and protein *Y*_*p*_ switches. The x-axis represents the number of switches that occurred during the simulation time interval, typically divided into discrete bins, while the y-axis shows the frequency of switches per trajectory. Due to noise, when the state switches from a lower state to a higher state, it is counted as one switch. The distribution of the number of switches in all trajectories across the complete time duration of the simulation is presented in Fig. 5. For the unconstrained scenario, where *α*_*p*_ = 80, the frequency of switches over 100 trajectories is shown in Fig. 5(d). In the constrained scenario, where *α*_*p*_ = 10, the number of switches per trajectory increased (Fig. 5(e)). Meanwhile, in the scenario where total ribosome *r*_*T*_ is varied alongside the translation rate *α*_*p*_ = 10, the switching frequency was substantially reduced as compared to the unconstrained scenario (Fig. 5(f)), making states more stable.

### B. Incoherent feedforward loop

In an incoherent feedforward loop (IFFL), an input signal regulates a target gene through two distinct pathways, direct activation and indirect repression, which oppose each other (Fig. 6(a), inset). In this incoherent feedforward loop, the input signal *u* directly activates the expression of both the genes *X* and *Y*, and simultaneously, the gene *X* encodes a repressor protein *X*_*p*_, which represses the gene *Y* . This regulatory configuration produces pulse-like dynamics [45].

**Fig. 6.**
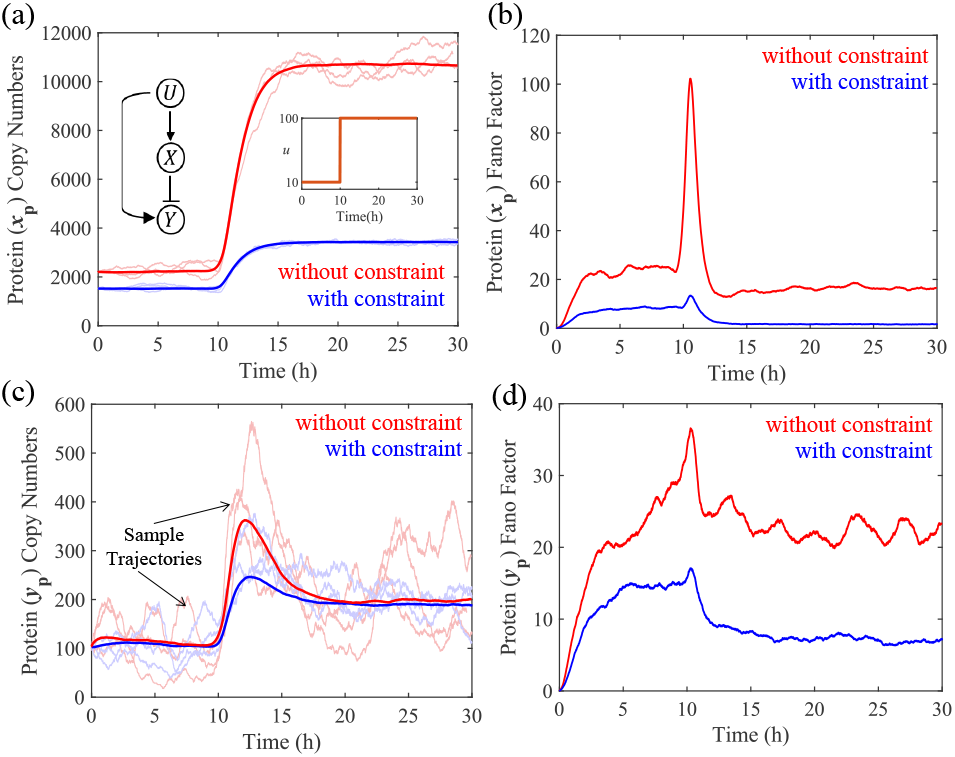
Simulations for incoherent feedforward loop for a step input of *u* from 10 to 100 at time = 10 *h* (inset, a) (Table VI). The key parameters for the model used were considered as follows: the rates are *α*_*mx*_ = *α*_*my*_ = 500 *h*^−1^, *γ*_*mx*_ = *γ*_*my*_ = 1 *h*^−1^, *k*_1_ = *k*_2_ = 100, *k*_*bx*_ = *k*_*by*_ = 1 *h*^−1^, *k*_*ux*_ = *k*_*uy*_ = 1 *h*^−1^, *γ*_*px*_ = *γ*_*py*_ = 1 *h*^−1^, and the total ribosome per cell *r*_*T*_ is given as 50 molecules. All these parameters are scaled with respect to the protein decay rate. (a) Mean of ***x***_*p*_ copy numbers, where blue solid line represents low translation rate *α*_*px*_ = *α*_*py*_ = 100 *h*^−1^ and red solid line represents high translation rate *α*_*px*_ = *α*_*py*_ = 1500 *h*^−1^, for 500 trajectories for protein *X*_*p*_. Sample trajectories for low resources and high resources are presented transparently in the respective colors, (b) Fano factor, variance over mean for 500 trajectories, of ***x***_*p*_ with low translation rate is represented as blue line and with high translation rate is represented as red line, (c) Mean of ***y***_*p*_ copy numbers, where blue solid line represents low translation rate *α*_*px*_ = *α*_*py*_ = 100 *h*^−1^ and red solid line represents high translation rate *α*_*px*_ = *α*_*py*_ = 1500 *h*^−1^, for 500 trajectories for protein *Y*_*p*_. Sample trajectories for low resources and high resources are presented transparently in the respective colors, (d) Fano factor, variance over mean for 500 trajectories, of ***y***_*p*_ with low translation rate is represented as blue line and with high translation rate is represented as red line.

**TABLE VI.**
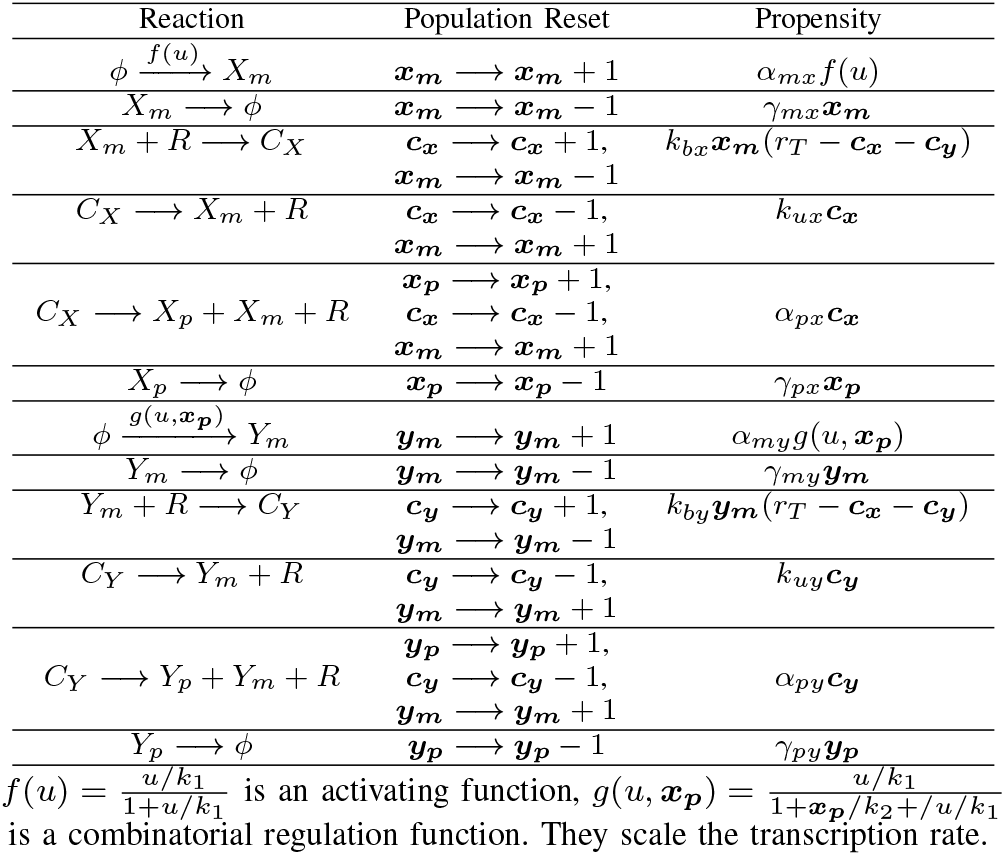
Propensities for population reset for corresponding reactions for Incoherent Feedforward Loop.

To model the incoherent feedforward loop using the stochastic simulation algorithm (SSA), we explicitly consider all relevant biochemical reactions. The probability per unit time for the reactions as a stochastic system can be written using propensity functions as stated in Table VI. The mRNA *X*_*m*_ is transcribed with the propensity *α*_*mx*_*f*(*u*) where 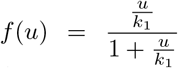 and *u* is a step signal and *k*_1_ is the dissociation constant, and this mRNA is degraded and diluted effectively with the propensity *γ*_*m*_***x***_*xm*_. The mRNA *X*_*m*_ binds to ribosomes *R* with the propensity *k*_*bx*_***x***_*m*_(*r*_*T*_ − ***c***_***x***_ − ***c***_***y***_) to form the translation complex *C*_*x*_, which dissociates with the propensity *k*_*ux*_***c***_*x*_. The translation complex *C*_*x*_ synthesizes protein *X*_*p*_ with the propensity *α*_*px*_***c***_*x*_, releasing free mRNA and ribosomes. The protein *X*_*p*_ undergoes dilution or degradation with the propensity *γ*_*px*_***x***_***p***_. Similarly, mRNA *Y*_*m*_ is transcribed with the propensity *α*_*my*_*g*(*u*, ***x***_***p***_), where 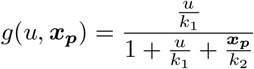, *k*_1_ is dissociation constant of *u*, and *k*_2_ is the dissociation constant of *X*_*p*_. The mRNA *Y*_*m*_ is degraded and diluted with the propensity *γ*_*my*_***y***_***m***_. The mRNA *Y*_*m*_ binds to ribosomes *R* with the propensity *k*_*by*_***y***_***m***_(*r*_*T*_ − ***c***_***x***_ − ***c***_***y***_) to form the translation complex *C*_*y*_, dissociating with the propensity *k*_*uy*_***c***_***y***_. This complex *C*_*y*_ produces protein *Y*_*p*_ with the propensity *α*_*py*_***c***_***y***_, again releasing free mRNA and ribosomes for reuse. Protein *Y*_*p*_ also undergoes dilution or degradation with the propensity *γ*_*py*_***y***_***p***_.

Similar to the single-gene system, resource availability can be modulated by varying the translation rate. At a high value of the translation rate, the number of free ribosomes in the cell is sufficient to approximate an unconstrained system. Conversely, at lower translation rates, resource availability drops significantly, introducing a constraint on gene expression within the IFFL.

The dynamics of protein concentration and the corresponding Fano factor for the incoherent feedforward loop are illustrated using stochastic simulations (Gillespie algorithm) in Fig. 6. The protein copy number ***x***_*p*_ is substantially reduced when the available resource is limited due to the low translation rate, compared to the scenario of a higher translation rate (Fig. 6(a)). This is expected as the limited resource creates a negative interaction among competitive species, as seen in single gene expression also. The Fano factor (variance-to-mean ratio) for protein *X*_*p*_ is notably lower under resource constraint conditions. Without constraints, the Fano factor stabilizes at a significantly higher value, indicating that the unconstrained scenario is more susceptible to stochastic variability (Fig. 6(b)). The presence of resource constraints reduces the amplitude of the protein pulse, indicating a direct impact of ribosomal limitation on the performance of the system (Fig. 6(c)). It is interesting to note that the resource constraint does not change the steady-state level or alter the adaptation capability of IFFL. Similar to the protein *X*_*p*_, with resource constraints, the fluctuations in protein *Y*_*p*_ are smaller, as evidenced by the lower Fano factor (Fig. 6(d)), reflecting reduced stochastic noise in the system. In both protein concentrations, lower variability and rapid adaptation suggest that resource constraints enhance robustness to noise.

## VI. CONCLUSION

In this work, we systematically investigated the effects of resource constraints on noise propagation in biomolecular systems. By comparing 3D, 2D, and 1D models of constitutive gene expression, we showed that protein-only models exhibit higher noise as compared to higher-order models with intermediate species. For single-gene expression, the Fano factor decreases with increasing resource constraint strength due to the saturating dependence of translation on ribosome availability. Extending to circuit motifs, we found that resource constraints reduce noise-induced switching frequency with fixed mean protein level in bistable toggle switches, enhancing robustness of bistability, while in incoherent feedforward loops, they lower the pulse amplitude and Fano factor without compromising adaptation. These results underscore the importance of incorporating both mechanistic detail and resource-aware considerations into the design of synthetic biomolecular circuits.

## VII. Future Work

In future work, we intend to add extrinsic noise and non-exponential timing of events [46], [47], to our resource-aware stochastic gene expression analysis.

## Appendix

### 1) Linearization of Nonlinear Systems

Consider a non-linear system:

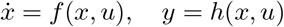

Let (*x*^∗^, *u*^∗^) be an equilibrium point. Defining deviations:

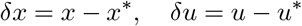

Performing Taylor expansion around equilibrium (*x*^∗^, *u*^∗^):

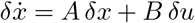

where,

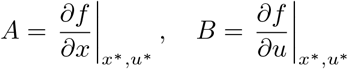

Similarly, the output equation becomes:

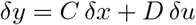

where,

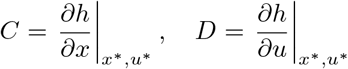

Thus, the linearized system is:

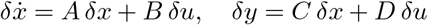

The complete expressions of the linearized matrices for all the models are:

3-dimensional model:

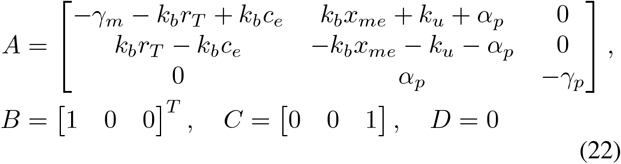

where, 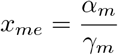 and 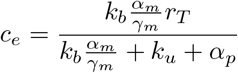.

2-dimensional model:

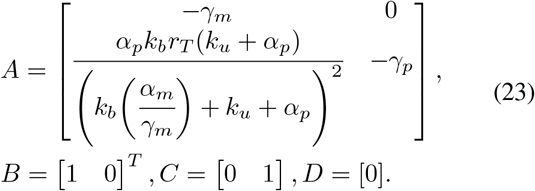

1-dimensional model:

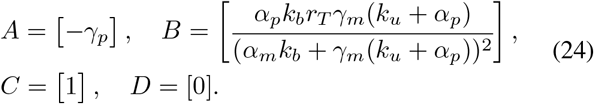

